# Correlated evolution of multiple traits gives butterflies a false head

**DOI:** 10.1101/2024.11.25.624978

**Authors:** Tarunkishwor Yumnam, Ullasa Kodandaramaiah

## Abstract

Many butterflies possess a combination of characters at the posterior hindwing end, superficially resembling their head. This ‘*false head*’ has been hypothesised to deflect predator attacks towards the false head area. A clear understanding of the diversity and evolution of false head traits across butterflies is lacking. Here, we tested whether false head traits evolved from simple to complex in order to achieve a greater resemblance to a head. We also tested if false head traits form an adaptive constellation and, thus, evolved correlatedly. Using a phylogenetic framework with 927 lycaenid species, our results illustrate evolutionary patterns of five false traits - (i) false antennae, (ii) spot, (iii) conspicuous colouration in the false head area, (iv) false head contour in the false head area, and (v) convergent lines. We found that false traits (i)–(iv) evolved in a correlated fashion across the phylogeny, likely driven by a common selective pressure. Our findings support the idea that false head functions as an adaptive constellation for predator attack deflection.

## Introduction

The enormous diversity in animal colouration has long fascinated humankind [1]. Although some animal colours are used for sexual signalling [2,3] and species recognition [4], many animals rely on colour patterns to protect themselves against predators. These protective strategies include various types of crypsis, which involve colour patterns to prevent detection or recognition by predators [5], aposematism, where bright colours advertise prey unprofitability [6], deimatism, wherein animals suddenly reveal their otherwise hidden conspicuous colours and cause a startle response [7], etc. Deflection is another anti-predator strategy that involves employing traits to manipulate the location on the prey’s body where the predator makes initial contact in a way that enhances the prey’s survival likelihood [8]. These deflective traits may bias the point of predator attack to the prey’s body parts that are less vulnerable and can be broken off, or away from the vital body parts. For example, many lizards have brightly coloured autotomous tails, which can direct avian attacks towards this expendable body part [9]. Longitudinal stripes on lizard bodies can also misdirect predator attacks towards the tail when the lizard is in motion [10]. One of the best-studied examples of deflective colour patterns is the eyespot, which is a “roughly circular pattern with at least two concentric rings or with a single colour disc and a central pupil” [11], potentially resembling the vertebrate eye. These eyespots occur across various taxa, including fish [12], frogs [13] and insects, particularly in the order Lepidoptera [14]. Small eyespots on the lepidopteran wing periphery may protect their bearers by deflecting attacks away from vital organs and towards the non-vital parts of the wings (reviewed in [15]). Marginal eyespots on the hindwing of the butterfly *Bicyclus anynana* have been shown to deflect mantid predation towards the less vulnerable hindwing area [16]. A deflective effect of eyespots has also been demonstrated in aquatic prey in the context of fish predation [17]. Moreover, dark and roughly circular spots, which may visually resemble a reduced eyespot, can provide a deflective benefit. For example, contrasting dark spots on the tadpoles tails can direct dragonfly attacks towards the tail [18], resulting in significantly higher survival than when attacks are on the body [19].

An eyespot can be considered an individual trait. While such traits can be effective on their own, multiple traits could function together for increased efficiency to deflect predator attacks. When multiple traits function together in synergy, they are referred to as adaptive constellations [20]. For example, many animals possess an eye stripe, which is a bar running through the eye, with its colour matching (part of) the eye. Such eye stripes in fishes have been suggested to impart a concealing effect on the eye [21,22]. A comparative study of butterflyfishes showed that all fish with eyespots also have eye stripes [12]. Through a series of predation experiments, Kjernsmo and Merilaita [17] first provided evidence for the deflective effect of fish eyespots and the concealing benefits of eye stripes. Their results also suggested that these two traits with different functions formed an adaptive constellation to produce a combined effect that increased the probability of deflection. Similarly, some swallowtail butterflies have hindwing tails and conspicuous colouration near the hindwing tail. Chotard and colleagues [23] showed that these two traits can function together to deflect bird attacks. They also demonstrated that the hindwing area near the tail was significantly easier to tear off compared to other parts of the wing, and thus enabled the butterfly to escape from a deflected attack. The tail, hindwing colouration and differential wing strength can be considered three traits that form an adaptive constellation to enhance the deflective effect.

A large number of butterflies possess a set of characters on the underside of their hindwings, popularly known as the false head due to the resemblance to the actual head of butterflies. Most butterflies rest with their wings closed, and, therefore, the false head is visible to predators when they are resting. The false head comprises a combination of traits: (i) tails, which presumably resemble false antennae, (ii) dark spots, (iii) conspicuous colouration, (iv) false head contour, and (v) convergent lines (Fig. 1; detailed definition in Methods). The false head is an example of a putative adaptive constellation that deflects predator attacks [24]. The false antennae, which move back and forth, are hypothesised to mimic the actual antennae, thus attracting attention towards themselves [24]. The quasi-circular dark spots of the false head can influence the initial contact point of predation [25]. The conspicuous colouration may help grab predators’ visual attention to the false head area [24,26]. Meanwhile, a modified contour of the posterior hind wing may provide a visual resemblance to the contour of a butterfly’s head [24]. Convergent lines may act as leading lines to direct the predator’s attention to the false head area [24,27].

**Figure 1.**
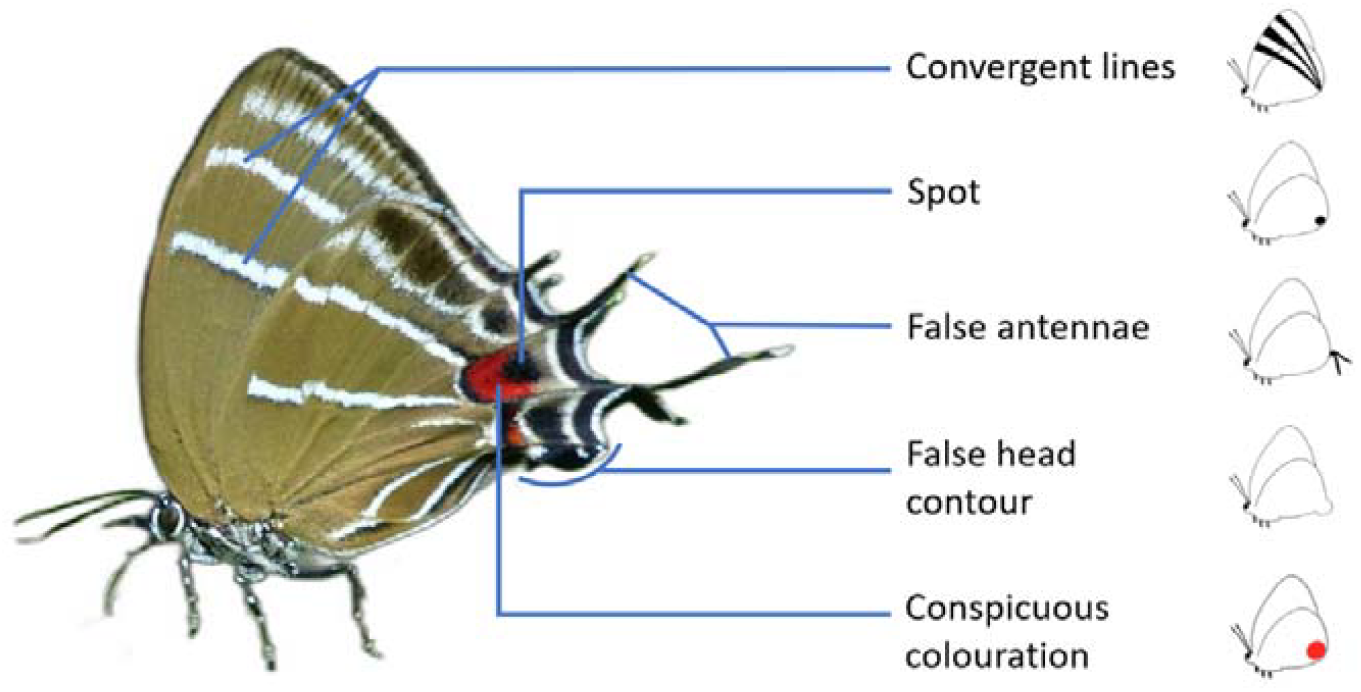
Photograph of *Airamanna columbia* depicting the five false head traits characterised in this study. Photo by Mark Eising (www.markeisingbirding.com): reproduced here with permission.

Compared to other body parts, physical damage in the false head area is less costly to the adult butterfly in terms of survival and behaviour, including flight dynamics [28] and mating [29]. From observations on wing damage in a large number of lycaenids, Robbins [30] argued that species with more false head traits were significantly more likely to deflect predator attacks successfully. Because attack deflection by false heads happens when butterflies are at rest, such failed attacks are expected to lead to symmetric damage on the wings, i.e., similar damage on the left and right wings [31]. A study based on pinned museum specimens revealed that species with more false head traits had higher symmetrical wing damage in the false head region, compared to species with fewer false head traits. This may be because having more complex false heads increases the probability of attacks being deflected [27]. Wourms and Wasserman [25] manually added false heads on the white wings of the *Pieris rapae* butterfly by painting and attaching multiple false head trait(s) singly and in combinations, and exposed these wings to birds (Great Tits – *Parus major*). Although only spots influenced the first attack contact point, the number of false head traits influenced prey handling location by the birds – models with more false head traits were more likely to be handled at the hind wing area compared to models with single or no false head traits. In contrast, López-Palafox & Cordero [32] did not find any difference in the mantid predation rates between hairstreak butterflies with intact and ablated false antennae, suggesting that, at least against some predator groups, a single false head trait independently may be ineffective in deflecting attacks.

Especially common in Lycaenidae and Riodinidae, some or all false head traits are also present in Papilionidae, Nymphalidae and Hesperiidae, i.e., in 5 of the 7 butterfly families. The presence of these false head traits varies across species, even within the same genus, as well as between sexes [27]. Thus, there is extensive variation in the morphology of false heads, but an understanding of the diversity and evolution of false head traits across butterflies is lacking. Here, we use phylogenetic comparative approaches to test hypotheses about the evolution of the false head. Specifically, we formulate hypotheses to test whether the false head complex functions as an adaptive constellation for attack deflection. If a false head works to deflect attacks towards a less vulnerable area, this less vulnerable area should be highly conspicuous in order to grab the predator’s attention. Moreover, if the false head deflects predator attacks because it fools the predators into perceiving it as a head, the false head should evolve to have many head-like characteristics. Thus, we predicted that the false head traits evolved from simple to complex, more elaborate, traits. We also reasoned that the false head traits function together as an adaptive constellation. Hence, we predicted that the false head traits evolved in a correlated pattern across the phylogeny.

It is possible that the body size of butterflies influences the evolution of the false head. Multiple comparative analyses have shown that body size can affect the evolution of body colour. Hossie and colleagues [33] demonstrated that the evolution of conspicuous eyespots in hawkmoth larvae was associated with larger body size. In the aposematic poison frogs, the evolution of larger body size was associated with increased conspicuous colouration. Moreover, the loss of conspicuous colouration in poison dart frogs likely co-evolved with a decrease in body size [34]. Similarly, reduced camouflage decoration behaviour in *Majoidea* crabs correlates with larger body sizes [35]. Here, we tested the evolutionary relationship between false head and wingspan. Because larger butterflies are easier to detect, they are more likely to rely on defences such as false heads. Therefore, we predicted that larger butterflies are more likely than smaller butterflies to have complex false heads. Alternatively, this prediction might be invalid if small and large butterflies face similar predation pressure from visually hunting predators despite having different predator types.

## Methods

We categorised false head traits as 5 discrete traits and collected data on the presence of these traits across 928 butterfly species by analysing images available in online databases (Supplementary: Table S1). We reconstructed the phylogeny of the species in this study using gene sequence data from NCBI (details in Methods: *Tree Building*) and performed multiple phylogenetic comparative analyses to test our hypotheses regarding the macroevolutionary pattern of false head evolution.

### Collection of false head data

The dataset on the presence and absence of the five false head traits – (i) false antennae, (ii) spot, (iii) conspicuous colour, (iv) false head contour and (v) convergent lines (Fig. 1) – was built as binary data with 1 representing presence and 0 absence. Data were coded from images of butterfly individuals with the false head region clearly visible and without wing damage. Images were accessed from online sources and databases (Supplementary: Table S1). We chose species for which at least one clear image of the undamaged underside of the butterfly was available. We relied on published descriptions with images and illustration plates for two species: *Arhopala tyrannus* (Taf. XXIXX, Fig. 1 & 2 in [36]) and Eirmocides ardosiacea (PIate I, Fig. 122-123, PIate 2, Fig. 133-134 in [37]). Whenever multiple images were available for a species, a minimum of two images were analysed. With these criteria, images representing 928 species were analysed by the authors, and the presence and absence of the false head traits were recorded as discrete traits. Some species exhibited sexual dimorphism in the presence of false head traits. In such cases, we considered data from the sex which possessed more false head traits. We first restricted our analysis to the family Lycaenidae, which has the highest number of species with false heads. The included species represented all major clades and broad regions of the distribution of Lycaenidae [38]. We also conducted similar analyses using a higher-level phylogeny [38] of 197 species, representing all families and 98% of butterfly subtribes. We acknowledge that colour patterns should ideally be assessed from images taken under controlled conditions. However, the scope of the study necessitated the availability of a large number of images from species from many countries across multiple continents. Therefore, it was not logistically possible for us to photograph all species under controlled conditions. Other studies with a similar scope as ours have employed human observers to grade animal colour patterns based on images [12,39–42].

We defined the false head area of a butterfly as the hindwing tornal area enclosed from above by the hindwing vein M3 and laterally by 50% of the hindwing inner margin. We defined the false head traits (Fig. 1) by modifying previously used definitions [26,27,30].

- **False antennae:** Thin antennae-like elongated protrusions in pair(s), regardless of the length and number of pairs, from the false head region of both hindwings were defined as false antennae.
- **Spot:** A spot was recorded present if a species had a quasi-circular dark spot in the false head area. However, if multiple such spots were present in other parts of the wing, a spot was not marked present (e.g., in the case of serial marginal eyespots as in *Mycalesis mineus*).
- **Conspicuous colour:** Conspicuous colour was recorded present if a species had a salient colour patch in the false head area, which (i) was absent in other wing areas, (ii) contrasted with the colours of the rest of the wings, and (iii) had a diameter larger than that of the spot.
- **False head contour:** False head contour was recorded present if the tornal wing area was modified by a protrusion of the wing shape, visually mimicking the contour of a butterfly head.
- **Convergent lines:** Convergent lines were defined as the presence of at least two lines running continuously across the ventral side of both wings and converging and meeting at the false head area.

### Tree building

Sequence data for nine genes, carbomylphosphate synthase domain protein (CAD) gene, cytochrome c oxidase subunit I (COI), elongation factor 1 alpha (EF1a), dopa decarboxylase (DDC), glyceraldehyde-3-phosphate dehydrogenase (GAPDH), histone H3 (H3), malate dehydrogenase (MDH), ribosomal protein S5 (RpS5) and wingless, were downloaded for 927 Lycaenidae and 1 Riodinidae species from Genbank [43] using the NSDPY package *v0*.*2*.*3* [44]. The riodinid served as the outgroup. Each gene dataset was aligned independently through the MAFFT web server (https://mafft.cbrc.jp/alignment/server/) with the default setup [45]. The aligned sequences were transformed into sequential nexus files using the ALTER web server [46]. Since some species had missing sequence data for particular genes, we adopted a constrained tree-building approach using a recently published phylogeny of butterflies [47] as the backbone. We reconstructed a maximum-likelihood phylogeny using the constrained tree search approach in IQTREE *v2*.*2*.*2*.*6* [48]. The constrained tree file was built by pruning a recent global butterfly phylogeny [47] and retaining species which were common with our species list with false head data. The best substitution model (GTR+F+I+R8) was selected using the *ModelFinder* option, and the best partition model, using the recommend *-p* command. Branch support of the tree was inferred using 1000 replicates of ultrafast bootstraps using *-B 1000* [49].

Time calibration of the tree was performed with MEGA *v11*.*0*.*13* [53] using the *RelTime-ML* approach [54] by choosing the settings suggested by Mello [55]. Secondary calibration points from [51] were implemented as minimum and maximum constraints with uniform distribution at six ingroup nodes of the crown groups representing Aphnaeinae (minimum: 34.4 million years ago (MYA) and maximum: 42.2 MYA), Curetinae (minimum: 5.4 MYA, maximum: 6.5 MYA), Lycaeninae (63.7 and: 65.4 MYA), Miletinae (57.1 and 58.9 MYA) and Poritiinae (41.1 and 43.5 MYA). The resulting ultrametric tree (Fig. 2) was used for further analyses.

**Figure 2.**
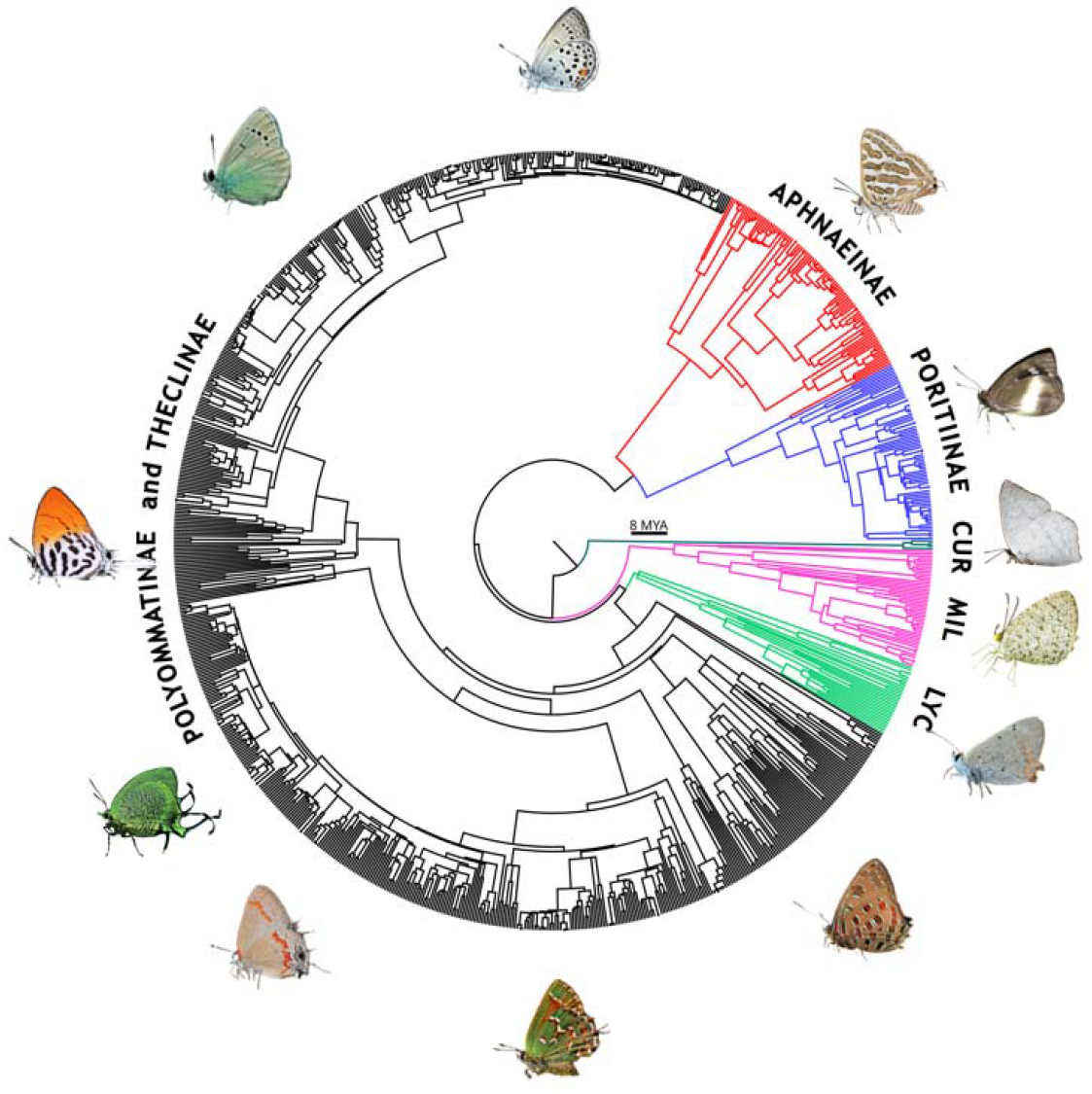
Maximum likelihood tree estimated in IQTREE using the constrained tree search approach and time-calibrated with RelTime-ML in MEGA. Clades belonging to a particular subfamily are coloured with the subfamily names written on the rim, where CUR: Curetinae, MIL: Miletinae and LYC: Lycaeninae. Photographs are reproduced from Wikimedia Commons, licensed under CC BY-SA with credits (clockwise from the TOP) to Ilia Ustyantsev, Gideon Pisanty, Charles J Sharp, Atanu Bose, zleng, Ivar Leidus, gailhampshire, khteWisconsin, Judy Gallagher, Kozue Kawakami, Cheongweei Gan and Hectonichus.

### Markov models for modelling false head evolution and ancestral state reconstruction

All analyses were conducted using the package *phytools v2*.*3*.*0* [50] in R *v4*.*2*.*1* [51] through Rstudio *v2023*.*03* [52]. We fitted equal rates (ER) and all rates different (ARD) time-continuous Markov (Mk) models to each false head trait. We used maximum likelihood to fit the models using *the fitMk* function, and the best-fitting model was selected based on the Akaike Information Criteria score. Furthermore, all the models were fitted with two different root priors. Models with a “flat” prior treated all states as having equal probabilities at the root [53], whereas those with a “fitzjohn” root prior regarded the root state as a nuisance parameter and implement an alternative root assignment by weighing each root state by its probability of giving rise to the extant data, considering the tree and model parameters [54]. Ancestral state reconstructions of the five false head traits were performed using Bayesian stochastic character mapping using the *make*.*simmap* function in the R package *phytools v2*.*3*.*0* [50] with 1000 simulations for both flat and fitzjohn root priors. The 1000 stochastic maps were summarised to calculate each state’s probability at the nodes, and the transition numbers were counted using the *countSimmap* function and averaged.

### Correlated evolutionary patterns between discrete traits

To analyse the correlated evolution of traits across the phylogeny, we used hidden Markov models implemented in the *corHMM package* [56]. For all possible pairwise combinations of the five false head traits, we used *fitCorrelationTest* function within *corHMM v2*.*8* to build four evolutionary models. We built two classic evolutionary models: Pagel’s independent model, which assumed that the two traits evolved independently of each other, and Pagel’s correlated model, which assumed the evolution of one trait was dependent on the other trait [57]. Hence, Pagel’s models had one rate category with transition rates between the four observed states of the pairwise binary traits. We also built hidden Markov independent model and correlated models. The former assumes no correlation, while the latter assumes a correlation between the focal traits [56]. In the hidden Markov models, we included two rate categories: one for the observed states and the other to account for unobserved conditions associated with the observed states, thus incorporating hidden evolutionary rate shifts. The hidden Markov models allowed heterogeneity in the transition rates between states. The best-fitting models were assessed based on the corrected AIC values. Adopting the same methods, we also tested the correlated evolutionary pattern of false head traits across all butterfly families using a pruned time-calibrated phylogeny derived from the phylogeny by Espeland and colleagues [38]. The pruned phylogeny had 192 species representing all butterfly subfamilies and subtribes except Pseudopontiinae. For taxa where only the genus was represented in the phylogeny, we coded a false head trait as present if any species within the genus had the particular false head trait.

### Phylogenetic path analyses to estimate the sequence of trait evolution

We inferred the probable order of evolution of the five false head traits using phylogenetic path analysis based on the d-separation method [58] using the R package *phylopath v1*.*1*.*3* [59] employing the phylogenetic linear regression approach and Pagel’s lambda evolution model. We considered all the possible path models (a total of 120) among the five false head traits. Best-fitting models were chosen based on the model ranking quantified using the C-statistics information criterion (CICc).

### Effect of wingspan on false head evolution

We used wingspan as a proxy for body size. We employed phylogenetic generalised least square regression using the R packages nlme *v3*.*1*.*164* [60] and *geiger v2*.*0*.*11* [61] to test the association between wingspan and false head evolution. We derived a dataset of 352 species for which we could obtain published wingspan data from the sources listed in Supplementary Table S1 and LepTraits 1.0 [62]. For species with minimum and maximum wingspan data, or sexual dimorphism in wingspan, we used the averaged measure for our analyses. We applied three evolutionary models (Ornstein–Uhlenbeck, Brownian motion and estimated lambda model) and relied on Akaike weights to choose the best-fitting model.

## Results

### Markov models for modelling false head evolution and ancestral state reconstruction

The ARD model was the best fitting across the models for all false head traits (Supplementary: Table S2). The average total number of transitions between the states remained similar across the two different root priors (Supplementary: Table S3). Similarly, the evolutionary transition rates between the presence and absence states of false head traits when fitted with fitzjohn root prior (Fig. 3(f)) did not differ from that of flat root prior (Supplementary: Fig. S1), and hence the results based on the fitzjohn prior are reported hereon. The ancestral state reconstruction of false head traits recovered multiple origins of false antennae (19.19 times), spots (43.62 times), conspicuous colouration (55.35 times), false head contour (10.12 times) and convergent lines (15.40 times) across the phylogeny. The analysis also recovered multiple losses of false antennae (83.64 times), spots (74.20 times), conspicuous colouration (88.34 times), false head contour (47.03 times) and convergent lines (13.33 times). The total number of character changes derived from models with flat root priors is given in Supplementary Table S3.

**Figure 3.**
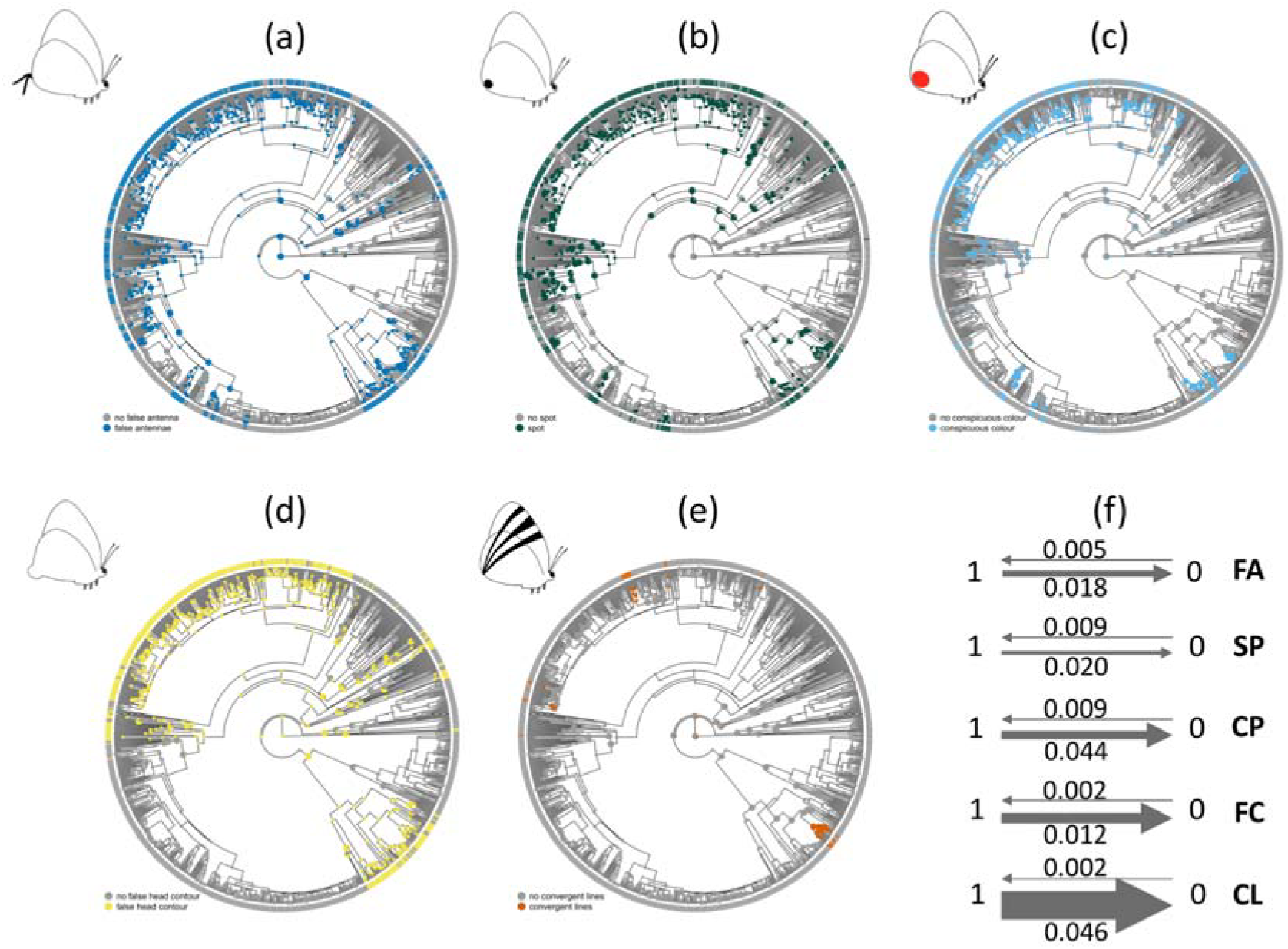
Ancestral state estimation using stochastic mapping with fitzjohn root prior for (a) false antennae, (b) spot, (c) conspicuous colouration, (d) false head contour and (e) convergent lines. The ring at the rim of the phylogeny represents the state of the extant taxa (grey for absence, coloured for presence). Pie charts represent the marginal frequencies of the states at the internal nodes. The size of the pies represents the probability for a state to occur at the nodes. Note that pies with a probability > 0.98 are reduced in size. (f) Evolutionary transition rates between absence (0) and presence (1) states of each false head trait derived from Mk models with fitzjohn root prior (FA: false antennae, SP: spot, CP: conspicuous colouration, FC: false head contour and CL: convergent lines). For each trait, the width of the arrows corresponds to the associated transition rates.

### Correlated evolutionary patterns between discrete traits

In our phylogenetic correlation analyses for all the possible pairwise traits, hidden Markov models were the best-fitting models. Hidden Markov correlated evolutionary models were favoured over all independent models and Pagel’s correlated models for all the possible pairwise combinations among false antennae, spots, conspicuous colouration and false head contour (Table 1). Therefore, the analyses indicate that these four traits evolved in a correlated fashion. The analyses also supported a correlated pattern of evolution of convergent lines with false head contour but not with the remaining false head traits (Table 1). To further investigate the evolutionary patterns of these traits at a broader phylogenetic scale, we conducted similar pairwise analyses using a time-calibrated phylogeny [38] representing all butterfly subfamilies and 98% of all butterfly subtribes. Here, Pagel’s correlated and independent models were favoured over hidden Markov models. The analyses indicated that false antennae, spot, conspicuous colouration and false head contour evolved in a correlated pattern, whereas convergent lines evolved independent of other false head traits (Fig. S2; Table S4).

**Table 1.**
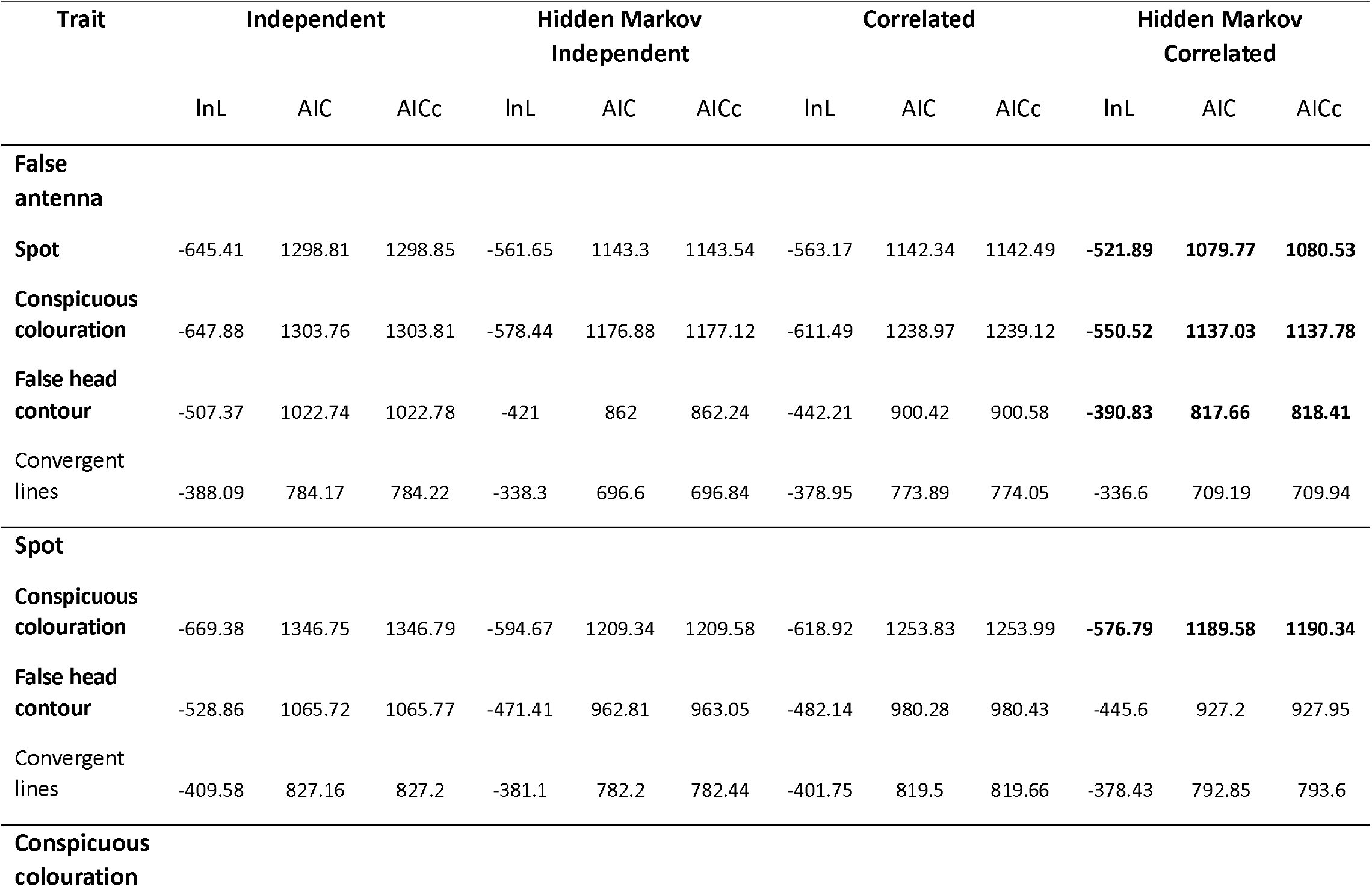
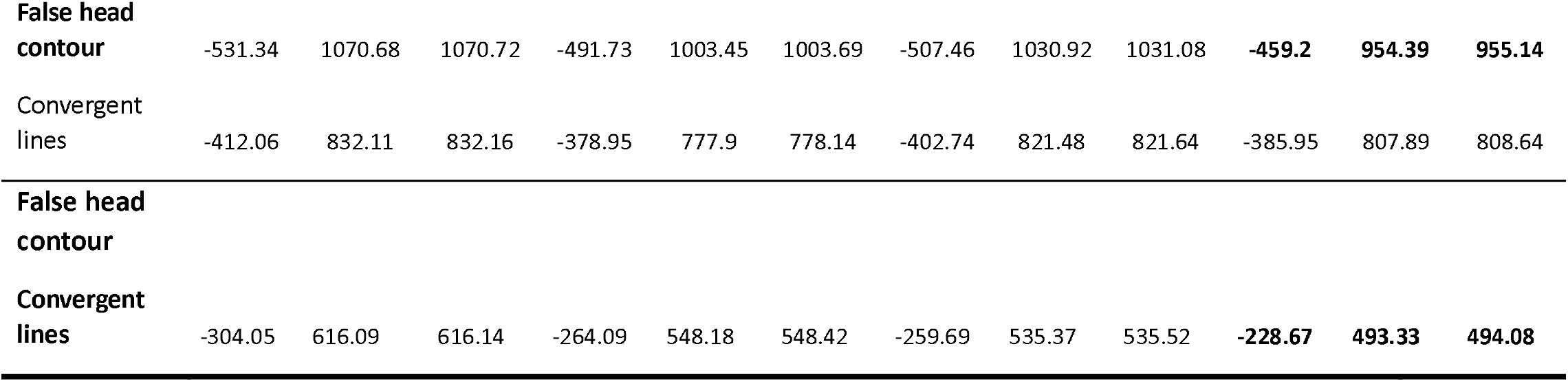
Evolutionary correlates of pairwise discrete false head traits resulted from corHMM analyses. Loglikelihood (lnL), AIC and corrected AIC or AICc of each evolutionary model are listed in the columns with the respective model’s name as the title. Model fit was assessed using the AICc. Pairwise traits with support for correlated evolution models are highlighted in bold.

### Phylogenetic path analyses to estimate the sequence of trait evolution

We built all the possible (total of 120) phylogenetic path models to explore the order of evolution of the five false head traits (Table2; Fig. 4). The best-fitting model (Table 2) indicated that conspicuous colouration positively influenced the evolution of false antennae (standard regression coefficient = 0.00002). Further, the evolution of spot is positively influenced by false antennae (standard regression coefficient = 0.427), false head contour by spot (standard regression coefficient = 0.166) and convergent lines by false head contour (standard regression coefficient = 0.114) (Fig. 4: RIGHT). Our result, thus, suggests that the probable direction of the evolution of traits is from conspicuous colouration to convergent lines via false antennae, spot and false head contour, in the respective order.

**Table 2.** Summary statistics of the 6 path models with the least CICc 502 from the phylogenetic path analysis, where C and ωCICc values indicate Fisher’s C-statistic and CICc weights, respectively. The models mentioned are as in Figure 4.

| Model | C | CICc | ΔCICc | likelihood | ωCICc |
| --- | --- | --- | --- | --- | --- |
| eighteen | 54.5 | 72.7 | 0.00 | 1 | 0.85 |
| ninety-five | 59.6 | 77.8 | 5.06 | 0.08 | 0.07 |
| ninety-six | 60.1 | 78.3 | 5.62 | 0.06 | 0.05 |
| two | 62.1 | 80.3 | 7.58 | 0.02 | 0.01 |
| thirty-six | 63.6 | 81.8 | 9.08 | 0.01 | 0.01 |
| sixteen | 73.8 | 92.0 | 19.31 | <0.01 | <0.01 |

**Figure 4.**
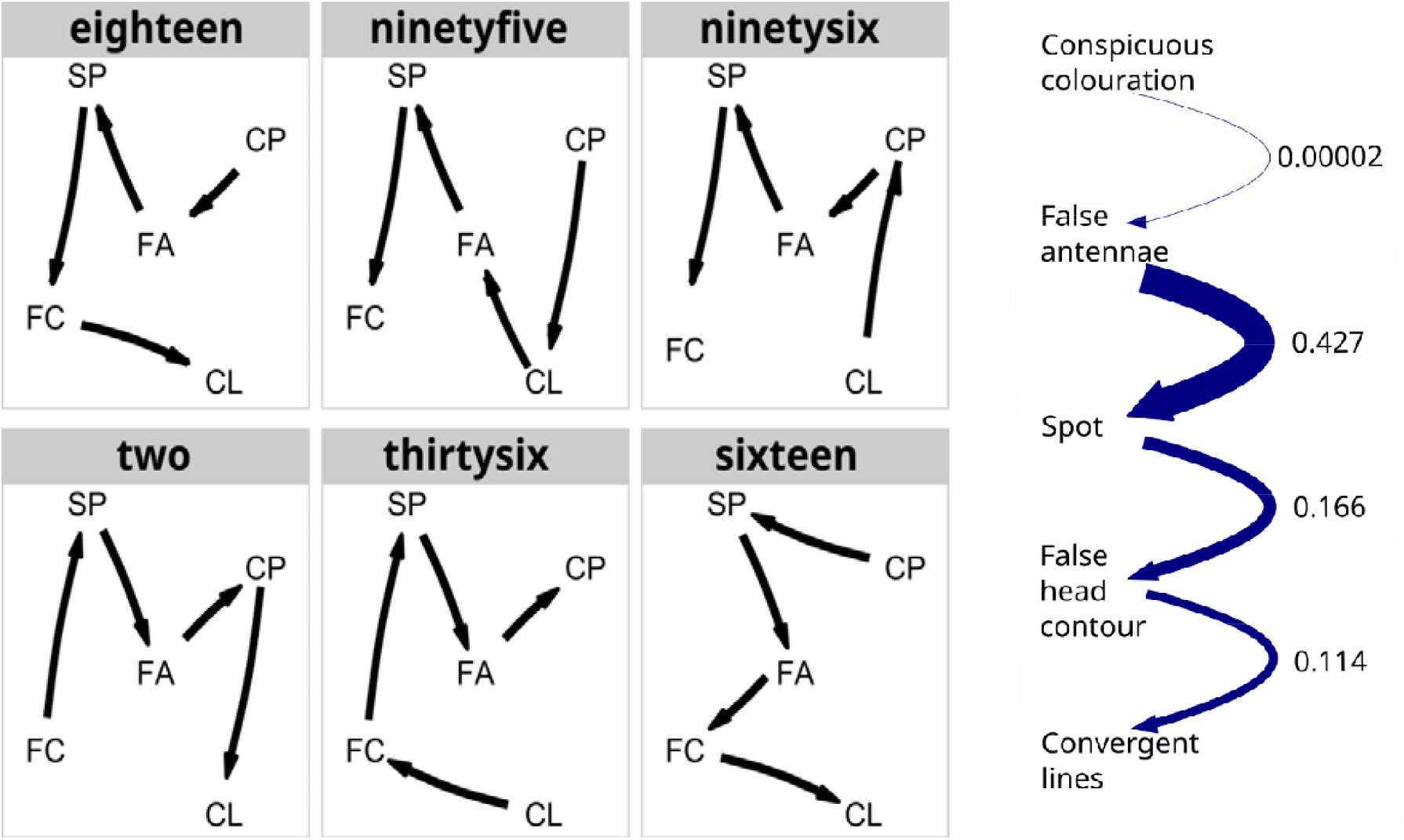
LEFT: Schematics of six (out of 120) phylogenetic path models having the least CICc values. Each model represents a hypothesised directionality among five false head traits: SP: spot, CP: conspicuous colouration, FA: false antennae, FC: false head contour and CL: convergent lines. Please refer to this figure in relation to Table 2. RIGHT: The best-fitting phylogenetic model suggests the evolutionary path starting with conspicuous colouration to false antennae, spot, and then false head contour and ending with convergent lines. The width of the curved paths corresponds to the standardised regression coefficients indicated on the right side of the curves. Note that the width of the curved path from conspicuous colouration to false antennae has been adjusted for visibility.

### Effect of wingspan on false head evolution

The phylogenetic generalised least square regression analysis tested the effect of the wingspan of butterflies on the evolution of false heads. The estimated lambda model (λ = 0.95, AIC = 483.81) was most favoured among the three evolutionary models (AIC of Ornstein–Uhlenbeck and Brownian motion were 660.66 and 3770.30 respectively). However, there was no phylogenetic association between an increase in wingspan and an increased number of false head traits (coefficient = 0.01, p = 0.644, Fig. 5), suggesting that wingspan did not influence the evolution of false head traits.

**Figure 5.**
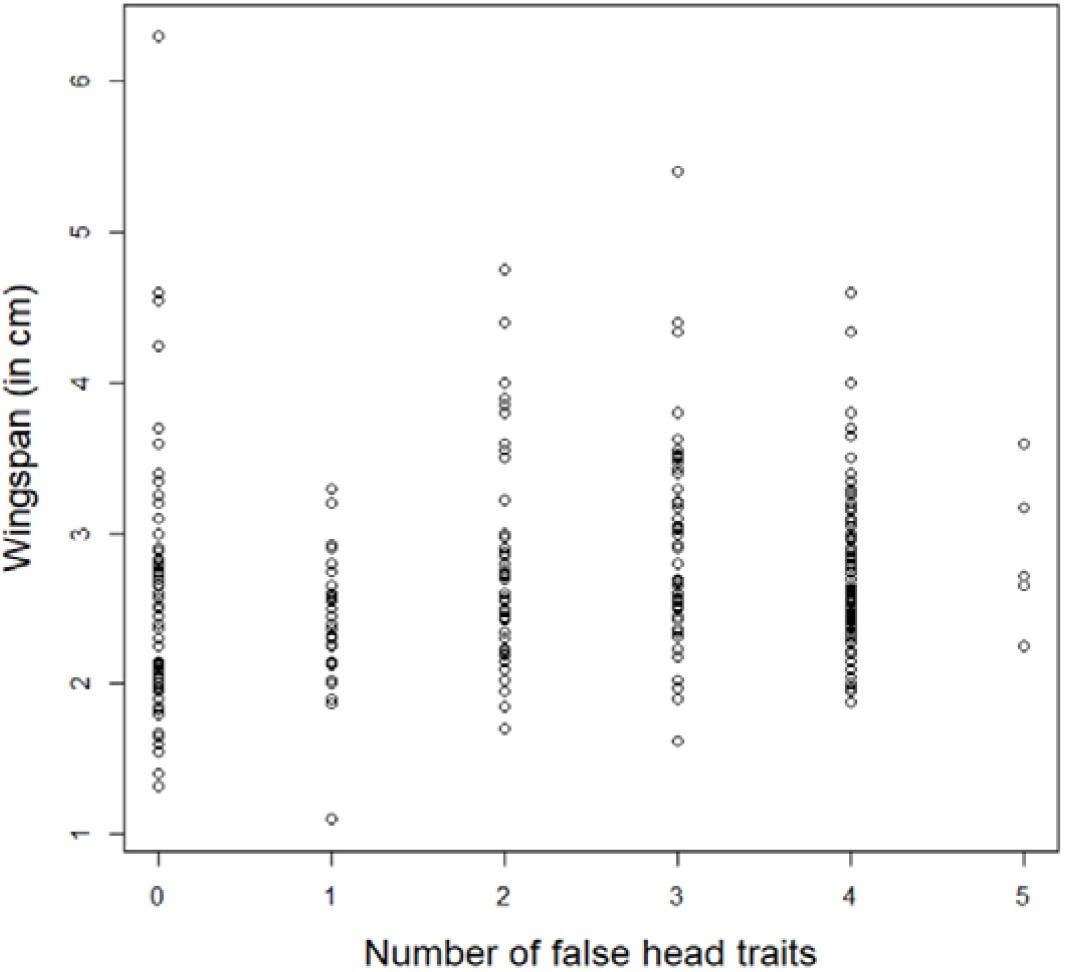
Plot showing no phylogenetic association between the number of false head traits and wingspan under the lambda model of evolution.

## Discussion

Despite several observational studies on the diversity and behaviour of false head traits, and experiments to understand their functional significance as anti-predator defences, an understanding of the evolution of the false head in a macroevolutionary context is lacking. Our phylogenetic comparative analyses using a large dataset of nearly 1000 lycaenid species shed light on the macroevolutionary trends of the evolution of various false head traits and indicate a correlated evolutionary pattern of the traits constituting the false head.

### Many false head traits are evolutionary labile

Our ancestral state analyses recovered multiple gains and losses of all false head traits suggesting that these traits are evolutionarily labile. The lability may be explained by differences in selection pressure on false heads. Natural predators of butterflies include birds [25,63], lizards [64], spiders [65] and mantids [32]. A false head may facilitate survival from predator attacks by a predator community, and the same false head may be ineffective, or may even increase the likelihood of detection, when predated upon by a different community. When a prey population experiences relaxed selection from predators, it can lead to the reduction or loss of morphological traits used for anti-predator defences [66]. Conversely, evolutionary changes in traits due to selection imposed in a specific environment over several hundreds of generations can be reversed within a few generations when the ancestral environment is reintroduced [67]. Thus, it is possible to lose and gain false head traits due to changes in the predator community. Similarly, a false head may be beneficial or detrimental depending on the habitat and changes in the habitat may select for or against false heads over evolutionary timescales.

It has been postulated that complex traits, once lost, are unlikely to be regained – popularly referred to as Dollo’s law [68]. In the past few decades, phylogenetic comparative analyses have reported many cases violating Dollo’s law [69]. These include studies demonstrating the regain of lost complex traits such as phasmid wings [70], anuran tympanic middle ear [71], frog mandibular teeth [72], squamate limbs [73], etc. The reversal of a trait after its loss could be due to the preservation of the molecular blueprint associated with the trait, even during the trait’s absence [74,75].

### Correlated evolutionary pattern of false head traits and its significance

The four false head traits - antennae, spot, false head contour and conspicuousness - shared a correlated pattern of evolution, suggesting a functional association among these correlated traits. Our results align with the hypothesised function of false heads as anti-predator strategies that deceive predators into misrecognising the true head. Studies based on observations of butterfly wing damage patterns in natural settings [30,78], museum specimens [27] and predation experiments [25,65,77] have provided evidence that the efficacy of false heads in deflecting attacks from predators such as birds and jumping spiders significantly increases when the butterfly (model) has a higher number of false head traits. This is corroborated by the macroevolutionary patterns in our study, which suggest that having a single false head trait may not provide an effective anti-predatory function. Therefore, our results support the idea that the false head is an adaptive constellation.

A hypothesis related to the deflective function of the false head is that the false head area should be easier to tear than the rest of the wing areas. If deflection works as an anti-predatory strategy, the wing area where attacks are deflected should be easily detachable so that the butterfly can escape [15]. Hill and Vaca [79] compared the wing tear weights between a conspicuous marginal patch and a homologous non-conspicuous patch across three species of *Pierella*. They found that the species with the conspicuous hindwing patch had lower wing tearing weights than those lacking the patch. Kodandaramaiah [15] posited that a comparison of wing tear force at the eyespotted location and other non-eyespotted locations across the wing surface of species with putative deflective eyespots could provide an understanding of the mechanism involved in deflection. Using dummy butterflies with actual wings of the swallowtail *Iphiclides podalirius* and wild-caught great tits as predators, Chotard and colleagues [23] found that the location of wing damage as a result of predation was significantly higher in the hindwing area and especially higher in the tail area when compared with other regions in both forewing and hindwing. When the force required to break the wing was compared across multiple locations on the wing surface, they found that hindwings, particularly the hindwing tails, were significantly easier to damage. Thus, there is some evidence that deflective colour patterns on butterfly wings are present in regions that easily tear. Similarly, Robbins [24] noted that the false head region on the hindwing of *Arawacus aetolus* broke off more easily compared to wing parts closer to the body. Studies on wing tear strengths of lycaenid butterflies with and without false head traits and at different parts of the wings are needed to test this idea in the context of false heads.

Our results do not support the anti-false head hypothesis, which posits that the bright, conspicuous colouration in the false head region *per se* directs attacks towards the colouration [24]. This hypothesis predicts that the evolution of the salient conspicuous colouration alone should provide effective anti-predator functionality. If this is the case, conspicuous colouration should have evolved independently of the other false head traits. Instead, the correlated evolutionary pattern of conspicuous colouration with other false head traits suggests that conspicuous colouration works in conjunction with other false head traits.

Our ancestral state reconstruction showed that convergent lines had multiple independent origins in two subfamilies – Aphnaeinae and Theclinae. Convergent lines have been hypothesised to lead predator’s attention towards the false head region where the lines converge [24]. This hypothesis of leading lines has received support from studies on the visual attention of human volunteers [80], and it is widely implemented as a principle in photography [81]. Under the premise that convergent lines and the remaining false head traits share the common function of deflecting attacks to the posterior hindwing end, we had predicted convergent lines to evolve correlatedly with other false head traits. However, we found that convergent lines showed no correlated evolutionary pattern with the rest of the false head traits, except false head contour. Hence, our results indicate that convergent lines do not share a functional association with the other traits. One possibility is that convergent lines, which often contrast strongly against the wing background colour, may act as disrupting colour patterns, as has been shown for similar patterns in the Banded Swallowtail (*Papilio demolion*) butterfly [82]. However, we do not disregard the possibility that the convergent lines evolved independently, providing a deflective function in a similar manner to the deflective effects of stripes on lizards, which redirect attacks towards the tail [10,83].

The combination of false head traits varies markedly across species, even within the same genera [27]. In our data, we recovered similar instances of multiple within-genus variations in the number and combination of false head traits. This variation may be explained by differences in selection pressures, rather than developmental or genetic constraints [30]. The differing numbers of false head traits likely represent alternative adaptive solutions to varying selection pressures from visual predators [27]. Furthermore, learning by predators may lead to frequency-dependent selection such that false heads confer higher fitness in butterflies communities where they are rare. Thus, predator learning may also contribute to inter-specific variation in false head traits.

Apart from their role in anti-predator functions, the presence of false head traits may incur energetic costs on the bearer. Some species have extremely long false antennae, similar in length or longer than half their wingspan, such as *Arcas cypria* (Theclinae), *Zeltus amasa* (Theclinae) and *Eooxylides tharis* (Theclinae). Such long false antennae may compromise flight performance. False heads may also be energetically costly in terms of the production of structural colours. It is possible that the deflective advantage provided by long false antennae outweighs the energetic costs of flight and colour production. False heads, or components of false heads such as conspicuous colouration, may also function as sexual signals, and the potential interplay between these two functions warrants further investigation.

In addition to Lycaenidae, false heads have been reported in Riodinidae [84,85] and Nymphalidae [78,86]. In addition to the nymphalids and riodinids, we found species having at least one false head trait in Papilionidae and Hesperiidae. Using a 197 species phylogenetic tree representing 98% of all butterfly subtribes [38], we performed similar analyses as for Lycaenidae in order to understand the evolutionary patterns of false head traits on a broader scale. In agreement with the patterns that we found within Lycaenidae, we recovered a correlated evolutionary pattern of spot, false antennae, conspicuous colouration, and false head contours, while convergent lines did not show a correlated evolutionary pattern with any of the other false head traits (Supplementary: Table S7, Fig. S2). These results further strengthen the interpretations of our study.

### Body size does not affect false head evolution

Predation risk is positively dependent on size, where larger-sized prey are more prone to detection, whereas smaller prey gain more protection by being cryptic (reviewed in [87]). If larger butterflies are more easily detected, they may be more dependent on deflection, whereas smaller butterflies rely on crypsis alone. Thus, we predicted the evolution of more false head traits in larger butterflies compared to smaller butterflies. However, our study did not find any phylogenetic correlation between the number of false head traits a butterfly had and its wingspan. Thus, false heads may deflect attacks irrespective of size, either as the primary defence [23] or as a late-acting defence when camouflage fails, as suggested by Novelo Galicia and colleagues [27]. Instead of prey body size, the crucial criteria for deflection may be the size of the deflective features. Deflection is likely to work when the deflective features have an optimal size below which the features are too small for detection and beyond which predators are not drawn to attack [14]. Alternatively, deflection may be successful if the false head traits, irrespective of the wingspan, are more conspicuous than (parts of) the real head, thereby offering a super-stimulus to the predators [88]. Visually oriented predators of butterflies, such as birds [25,63], lizards [64], spiders [65] and mantids [32], may exert similar predation pressures on butterflies of varying sizes, driving the evolution of similar deflective features in varying prey sizes.

## Summary and Conclusion

The false head in butterflies has long been hypothesised to have a deflective function against predators. We present the first phylogenetic comparative analysis of false heads. We found that most false head traits in butterflies evolved in a correlated pattern, presumably towards a functional association as a response to a common selective force. Thus, our study provides macroevolutionary support for the idea that the false head evolved as an adaptive constellation of anti-predatory traits. These false head traits together may form a complex structure that visually resembles a head to the predators. Our results are in agreement with studies that showed that false heads deflect predator attacks efficiently when the butterfly has many false head traits. However, we also argue that multiple anti-predator hypotheses may work together synergistically along with the deflection hypothesis highlighting the importance of further predation experiments employing multiple predator types. Overall, our comprehensive analysis underscores further studies to understand the adaptive significance of false heads in butterfly defence mechanisms.

## Ethics

Our study does not involve any live animals and requires no ethical permits.

## Acknowledgement

We thank the anonymous Associate Editor and the two reviewers for their insightful comments, which improved our manuscript. We would like to thank (i) Vivek Philip Cyriac and Gopal Murali, IISc Bengaluru, for their suggestions in conducting the analyses; (ii) James Boyko, University of Michigan, for suggestions with corHMM; (iii) Mark Eising from The Netherlands (www.markeisingbirding.com), Fahim Khan, Ramón, Baranyi Tamas, Fahim Khan, mr_fab, steveball, portiod, adam187, Ken-ichi Ueda, Edward Perry, juancarlosgarciamorales1 for their butterfly images; (iii) Deepthi Padiyar for help in butterfly illustration and (iv) Akhil Sadiq, Indukala Prasannakumar, Anaswara K Sreedharan, Bhanu Bhakta Sharma, Shreya Mishra and Kushankur Bhattacharyya for their comments on the manuscript.

## References

1. Aristotle. 1910 Historia Animalium. Oxford: Translation by D’Arcy Wentworth Thompson. Clarendon Press. (https://archive.org/details/thompson-1910-aristotle-animals/mode/2up)

2. Emberts Z, Wiens JJ. 2022 Why are animals conspicuously colored? Evolution of sexual versus warning signals in land vertebrates. Evolution 76, 2879–2892. (doi:10.1111/evo.14636)

3. Svensson PA, Wong BBM. 2011 Carotenoid-based signals in behavioural ecology: a review. Behaviour 148, 131–189. (doi:10.1163/000579510X548673)

4. Klomp DA, Stuart-Fox D, Cassidy EJ, Ahmad N, Ord TJ. 2017 Color pattern facilitates species recognition but not signal detection: a field test using robots. Behavioral Ecology 28, 597–606. (doi:10.1093/beheco/arw186)

5. Stevens M, Merilaita S. 2009 Animal camouflage: current issues and new perspectives. Philos Trans R Soc Lond B Biol Sci 364, 423–427. (doi:10.1098/rstb.2008.0217)

6. Ruxton GD, Allen WL, Sherratt TN, Speed MP, Ruxton GD, Allen WL, Sherratt TN, Speed MP. 2018 Avoiding Attack: The Evolutionary Ecology of Crypsis, Aposematism, and Mimicry. Second Edition, pp 84–102. Oxford, New York: Oxford University Press.

7. Umbers KDL, De Bona S, White TE, Lehtonen J, Mappes J, Endler JA. 2017 Deimatism: a neglected component of antipredator defence. Biol Lett 13, 20160936. (doi:10.1098/rsbl.2016.0936)

8. Humphreys RK, Ruxton GD. 2018 What is known and what is not yet known about deflection of the point of a predator’s attack. Biological Journal of The Linnean Society 123, 483–495. (doi:10.1093/biolinnean/blx164)

9. Watson CM, Roelke CE, Pasichnyk PN, Cox CL. 2012 The fitness consequences of the autotomous blue tail in lizards: an empirical test of predator response using clay models. Zoology 115, 339–344. (doi:10.1016/j.zool.2012.04.001)

10. Murali G, Kodandaramaiah U. 2016 Deceived by stripes: conspicuous patterning on vital anterior body parts can redirect predatory strikes to expendable posterior organs. Royal Society Open Science 3, 160057. (doi:10.1098/rsos.160057)

11. Monteiro A. 2008 Alternative models for the evolution of eyespots and of serial homology on lepidopteran wings. BioEssays 30, 358–366. (doi:10.1002/bies.20733)

12. Kelley JL, Fitzpatrick JL, Merilaita S. 2013 Spots and stripes: ecology and colour pattern evolution in butterflyfishes. Proceedings of the Royal Society B: Biological Sciences 280, 20122730. (doi:10.1098/rspb.2012.2730)

13. Hernández-Palma TL, Rueda-Solano LA, Valkonen JK, Rojas B. 2023 Predator response to the coloured eyespots and defensive posture of Colombian four-eyed frogs. Journal of Evolutionary Biology 36, 1040–1049. (doi:10.1111/jeb.14193)

14. Stevens M. 2005 The role of eyespots as anti-predator mechanisms, principally demonstrated in the Lepidoptera. Biological Reviews 80, 573–588. (doi:10.1017/s1464793105006810)

15. Kodandaramaiah U. 2011 The evolutionary significance of butterfly eyespots. Behavioral Ecology 22, 1264–1271. (doi:10.1093/beheco/arr123)

16. Prudic KL, Stoehr AM, Wasik BR, Monteiro A. 2015 Eyespots deflect predator attack increasing fitness and promoting the evolution of phenotypic plasticity. Proceedings of the Royal Society B: Biological Sciences 282, 20141531. (doi:10.1098/rspb.2014.1531)

17. Kjernsmo K, Merilaita S. 2013 Eyespots divert attacks by fish. Proceedings of The Royal Society B: Biological Sciences 280, 20131458. (doi:10.1098/rspb.2013.1458)

18. van Buskirk J, Aschwanden J, Buckelmüller I, Reolon S, Rüttiman S. 2004 Bold Tail Coloration Protects Tadpoles from Dragonfly Strikes. Copeia 2004, 599–602. (doi:10.1643/CE-03-283R)

19. van Buskirk J, Anderwald P, Lüpold S, Reinhardt L, Schuler H. 2003 The Lure Effect, Tadpole Tail Shape, and the Target of Dragonfly Strikes. Journal of Herpetology 37, 420–424. (doi:10.1670/0022-1511(2003)037[0420:TLETTS]2.0.CO;2)

20. Kjernsmo K, Grönholm M, Merilaita S. 2016 Adaptive constellations of protective marks: eyespots, eye stripes and diversion of attacks by fish. Animal Behaviour 111, 189–195. (doi:10.1016/j.anbehav.2015.10.028)

21. Barlow GW. 1972 The Attitude of Fish Eye-Lines in Relation to Body Shape and to Stripes and Bars. Copeia 1972, 4–12. (doi:10.2307/1442777)

22. Neudecker S. 1989 Eye camouflage and false eyespots: chaetodontid responses to predators. Environ Biol Fish 25, 143–157. (doi:10.1007/BF00002208)

23. Chotard A, Ledamoisel J, Decamps T, Herrel A, Chaine AS, Llaurens V, Debat V. 2022 Evidence of attack deflection suggests adaptive evolution of wing tails in butterflies. Proceedings of the Royal Society B: Biological Sciences 289, 20220562. (doi:10.1098/rspb.2022.0562)

24. Robbins RK. 1980 The lycaenid false head hypothesis: historical review and quantitative analysis. Journal of the Lepidopterists’ Society 34, 194–208.

25. Wourms MK, Wasserman FE. 1985 Butterfly wing markings are more advantageous during handling than during the initial strike of an avian predator. Evolution 39, 845–851. (doi:10.1111/j.1558-5646.1985.tb00426.x)

26. Hendrick LK, Somjee U, Rubin JJ, Kawahara AY. 2022 A Review of False Heads in Lycaenid Butterflies. Journal of the Lepidopterists’ Society 76, 140–148. (doi:10.18473/lepi.76i2.a6)

27. Novelo Galicia E, Luis Martínez MA, Cordero C. 2019 False head complexity and evidence of predator attacks in male and female hairstreak butterflies (Lepidoptera: Theclinae: Eumaeini) from Mexico. PeerJ (doi:10.7717/peerj.7143)

28. Le Roy C, Cornette R, Llaurens V, Debat V. 2019 Effects of natural wing damage on flight performance in Morpho butterflies: what can it tell us about wing shape evolution? Journal of Experimental Biology 222, jeb204057. (doi:10.1242/jeb.204057)

29. Medina C, Cordero C. 2021 False heads and sexual behaviour in a hairstreak butterfly, Callophrys xami (Lepidoptera: Lycaenidae). European Journal of Entomology 118, 394–398. (doi:10.14411/eje.2021.040)

30. Robbins RK. 1981 The false head hypothesis: predation and wing pattern variation of lycaenid butterflies. The American Naturalist 118, 770–775. (doi:10.1086/283868)

31. Molleman F et al. 2020 Quantifying the effects of species traits on predation risk in nature: A comparative study of butterfly wing damage. Journal of Animal Ecology 89, 716–729. (doi:10.1111/1365-2656.13139)

32. López-Palafox TG, Cordero C. 2017 Two-headed butterfly vs. mantis: do false antennae matter? PeerJ (doi:10.7717/peerj.3493)

33. Hossie TJ, Skelhorn J, Breinholt JW, Kawahara AY, Sherratt TN. 2015 Body size affects the evolution of eyespots in caterpillars. Proceedings of the National Academy of Sciences 112, 6664–6669. (doi:10.1073/pnas.1415121112)

34. Rudh A. 2013 Loss of conspicuous coloration has co-evolved with decreased body size in populations of poison dart frogs. Evol Ecol 27, 755–767. (doi:10.1007/s10682-013-9649-8)

35. Hultgren KM, Stachowicz JJ. 2009 Evolution of Decoration in Majoid Crabs: A Comparative Phylogenetic Analysis of the Role of Body Size and Alternative Defensive Strategies. The American Naturalist 173, 566–578. (doi:10.1086/597797)

36. Wüllerstorf-Urbair B, Expedition N. 1861 Reise der österreichischen Fregatte Novara um die Erde in den Jahren 1857, 1858, 1859 unter den Befehlen des Commodore B. von Wüllerstorf-Urbair. Wien: Kaiserlich-Königliche Hof-und Staatsdruckerei; in Commission bei K. Gerold’s Sohn. (doi:10.5962/bhl.title.1597)

37. Tite GE. 1964 A Revision of the Genus Candalides. In Bulletin of the British Museum (Natural History): Entomology Vol. XIV, pp. 197–260. London: British Museum (Natural History).

38. Espeland M et al. 2018 A Comprehensive and Dated Phylogenomic Analysis of Butterflies. Current Biology 28, 770–778.e5. (doi:10.1016/j.cub.2018.01.061)

39. Allen WL, Baddeley R, Scott-Samuel NE, Cuthill IC. 2013 The evolution and function of pattern diversity in snakes. Behavioral Ecology 24, 1237–1250. (doi:10.1093/beheco/art058)

40. Arbuckle K, Speed MP. 2015 Antipredator defenses predict diversification rates. Proceedings of the National Academy of Sciences 112, 13597–13602. (doi:10.1073/pnas.1509811112)

41. Kodandaramaiah U. 2009 Eyespot evolution: phylogenetic insights from Junonia and related butterfly genera (Nymphalidae: Junoniini). Evolution & Development 11, 489–497. (doi:10.1111/j.1525-142X.2009.00357.x)

42. Murali G, Kodandaramaiah U. 2018 Body size and evolution of motion dazzle coloration in lizards. Behavioral Ecology 29, 79–86. (doi:10.1093/beheco/arx128)

43. Sayers EW, Cavanaugh M, Clark K, Pruitt KD, Schoch CL, Sherry ST, Karsch-Mizrachi I. 2021 GenBank. Nucleic Acids Research 49, D92–D96. (doi:10.1093/nar/gkaa1023)

44. Hebert R, Meglécz E. 2022 NSDPY: A python package to download DNA sequences from NCBI. SoftwareX 18, 101038. (doi:10.1016/j.softx.2022.101038)

45. Katoh K, Misawa K, Kuma K, Miyata T. 2002 MAFFT: a novel method for rapid multiple sequence alignment based on fast Fourier transform. Nucleic Acids Research 30, 3059–3066. (doi:10.1093/nar/gkf436)

46. Glez-Peña D, Gómez-Blanco D, Reboiro-Jato M, Fdez-Riverola F, Posada D. 2010 ALTER: program-oriented conversion of DNA and protein alignments. Nucleic Acids Research 38, W14–W18. (doi:10.1093/nar/gkq321)

47. Kawahara AY et al. 2023 A global phylogeny of butterflies reveals their evolutionary history, ancestral hosts and biogeographic origins. Nat Ecol Evol 7, 903–913. (doi:10.1038/s41559-023-02041-9)

48. Minh BQ, Schmidt HA, Chernomor O, Schrempf D, Woodhams MD, von Haeseler A, Lanfear R. 2020 IQ-TREE 2: New Models and Efficient Methods for Phylogenetic Inference in the Genomic Era. Molecular Biology and Evolution 37, 1530–1534. (doi:10.1093/molbev/msaa015)

49. Hoang DT, Chernomor O, von Haeseler A, Minh BQ, Vinh LS. 2018 UFBoot2: Improving the Ultrafast Bootstrap Approximation. Molecular Biology and Evolution 35, 518–522. (doi:10.1093/molbev/msx281)

50. Revell LJ. 2024 phytools 2.0: an updated R ecosystem for phylogenetic comparative methods (and other things). PeerJ 12, e16505. (doi:10.7717/peerj.16505)

51. R Core Team. 2023 R: A language and environment for statistical computing. R Foundation for Statistical Computing, Vienna, Austria. (https://www.R-project.org/)

52. RStudio Team. 2023 RStudio: Integrated Development for R. Rstudio, PBC, Boston, MA. (http://www.rstudio.com/)

53. Schluter D, Price T, Mooers AØ, Ludwig D. 1997 Likelihood of ancestor states in adaptive radiation. Evolution 51, 1699–1711. (doi:10.1111/j.1558-5646.1997.tb05095.x)

54. FitzJohn RG, Maddison WP, Otto SP. 2009 Estimating Trait-Dependent Speciation and Extinction Rates from Incompletely Resolved Phylogenies. Systematic Biology 58, 595–611. (doi:10.1093/sysbio/syp067)

55. Revell LJ. 2012 phytools: an R package for phylogenetic comparative biology (and other things). Methods in Ecology and Evolution 3, 217–223. (doi:10.1111/j.2041-210X.2011.00169.x)

56. Boyko JD, Beaulieu JM. 2023 Reducing the Biases in False Correlations Between Discrete Characters. Systematic Biology 72, 476–488. (doi:10.1093/sysbio/syac066)

57. Pagel M. 1997 Detecting correlated evolution on phylogenies: a general method for the comparative analysis of discrete characters. Proceedings of the Royal Society of London. Series B: Biological Sciences 255, 37–45. (doi:10.1098/rspb.1994.0006)

58. Hardenberg A von, Gonzalez-Voyer A. 2013 Disentangling evolutionary cause-effect relationships with phylogenetic confirmatory path analysis. Evolution 67, 378–387. (doi:10.1111/j.1558-5646.2012.01790.x)

59. Bijl W van der. 2018 phylopath: Easy phylogenetic path analysis in R. PeerJ 6, e4718. (doi:10.7717/peerj.4718)

60. Pinheiro J, Bates D, R Core Team. 2023 nlme: Linear and Nonlinear Mixed Effects Models. R package version 3.1.165. (https://cran.r-project.org/web/packages/nlme/index.html)

61. Harmon LJ, Weir JT, Brock CD, Glor RE, Challenger W. 2008 GEIGER: investigating evolutionary radiations. Bioinformatics 24, 129–131. (doi:10.1093/bioinformatics/btm538)

62. Shirey V et al. 2022 LepTraits 1.0 A globally comprehensive dataset of butterfly traits. Sci Data 9, 382. (doi:10.1038/s41597-022-01473-5)

63. Pinheiro CEG, Cintra R. 2017 Butterfly Predators in the Neotropics: Which Birds are Involved? Journal of the Lepidopterists’ Society 71, 109–114. (doi:10.18473/lepi.71i2.a5)

64. Halali D, Krishna A, Kodandaramaiah U, Molleman F. 2019 Lizards as Predators of Butterflies: Shape of Wing Damage and Effects of Eyespots. Journal of the Lepidopterists’ Society 73, 78–86. (doi:10.18473/lepi.73i2.a2)

65. Sourakov A. 2013 Two heads are better than one: false head allows Calycopis cecrops (Lycaenidae) to escape predation by a Jumping Spider, Phidippus pulcherrimus (Salticidae). Journal of Natural History 47, 1047–1054. (doi:10.1080/00222933.2012.759288)

66. Blumstein DT. 2006 The Multipredator Hypothesis and the Evolutionary Persistence of Antipredator Behavior. Ethology 112, 209–217. (doi:10.1111/j.1439-0310.2006.01209.x)

67. Teotónio H, Rose MR. 2000 Variation in the reversibility of evolution. Nature 408, 463–466. (doi:10.1038/35044070)

68. Gould SJ. 1970 Dollo on Dollo’s law: Irreversibility and the status of evolutionary laws. J Hist Biol 3, 189–212. (doi:10.1007/BF00137351)

69. Collin R, Miglietta MP. 2008 Reversing opinions on Dollo’s Law. Trends in Ecology & Evolution 23, 602–609. (doi:10.1016/j.tree.2008.06.013)

70. Forni G, Martelossi J, Valero P, Hennemann FH, Conle O, Luchetti A, Mantovani B. 2022 Macroevolutionary Analyses Provide New Evidence of Phasmid Wings Evolution as a Reversible Process. Systematic Biology 71, 1471–1486. (doi:10.1093/sysbio/syac038)

71. Pereyra MO et al. 2016 The complex evolutionary history of the tympanic middle ear in frogs and toads (Anura). Sci Rep 6, 34130. (doi:10.1038/srep34130)

72. Wiens JJ. 2011 Re-evolution of lost mandibular teeth in frogs after more than 200 million years, and re-evaluating Dollo’s law. Evolution 65, 1283–1296. (doi:10.1111/j.1558-5646.2011.01221.x)

73. Kohlsdorf T, Wagner GP. 2006 Evidence for the Reversibility of Digit Loss: A Phylogenetic Study of Limb Evolution in Bachia (Gymnophthalmidae: Squamata). Evolution 60, 1896–1912. (doi:10.1111/j.0014-3820.2006.tb00533.x)

74. Leal F, Cohn MJ. 2016 Loss and Re-emergence of Legs in Snakes by Modular Evolution of Sonic hedgehog and HOXD Enhancers. Current Biology 26, 2966–2973. (doi:10.1016/j.cub.2016.09.020)

75. Whiting MF, Bradler S, Maxwell T. 2003 Loss and recovery of wings in stick insects. Nature 421, 264–267. (doi:10.1038/nature01313)

76. Fordyce JA. 2006 The evolutionary consequences of ecological interactions mediated through phenotypic plasticity. Journal of Experimental Biology 209, 2377–2383. (doi: 10.1242/jeb.02271)

77. Bartos M, Minias P, Minias P. 2016 Visual cues used in directing predatory strikes by the jumping spider Yllenus arenarius (Araneae, Salticidae). Animal Behaviour 120, 51–59. (doi:10.1016/j.anbehav.2016.07.021)

78. Tonner M, Novotny V, Lepš J, Komarek S. 1993 False head wing pattern of the burmese junglequeen butterfly and the deception of avian predators. Biotropica 25, 474–478. (doi:10.2307/2388871)

79. Hill RI, Vaca JF. 2004 Differential wing strength in Pierella butterflies (Nymphalidae, Satyrinae) supports the deflection hypothesis. Biotropica 36, 362–370. (doi:10.1646/03191)

80. Mochizuki I, Toyoura M, Mao X. 2018 Visual attention prediction for images with leading line structure. Vis Comput 34, 1031–1041. (doi:10.1007/s00371-018-1518-6)

81. Zhang J, Synave R, Delepoulle S, Cozot R. 2024 Reconstructing Image Composition: Computation of Leading Lines. Journal of Imaging 10, 5. (doi:10.3390/jimaging10010005)

82. Tan EJ, Wilts BD, Tan BTK, Monteiro A. 2020 What’s in a band? The function of the color and banding pattern of the Banded Swallowtail. Ecology and Evolution 10, 2021–2029. (doi:10.1002/ece3.6034)

83. Murali G, Merilaita S, Kodandaramaiah U. 2018 Grab my tail: evolution of dazzle stripes and colourful tails in lizards. Journal of Evolutionary Biology 31, 1675–1688. (doi:10.1111/jeb.13364)

84. Robbins RK. 1985 Independent evolution of false head behavior in Riodinidae. Journal of The Lepidopterists’ Society 39, 224–225.

85. DeVries PJ. 1997 The Butterflies of Costa Rica and Their Natural History, Volume II: Riodinidae. First Edition. Princeton, New Jersey: Princeton University Press.

86. Sourakov A. 2015 Antipredation and ‘antimimicry’: wing pattern is supported by behavior in Archaeoprepona chromus (Lepidoptera: Nymphalidae: Preponini). Association for Tropical Lepidoptera Notes 2015, 1–7.

87. Yu H, Lin Z, Xiao F. 2024 Role of body size and shape in animal camouflage. Ecology and Evolution 14, e11434. (doi:10.1002/ece3.11434)

88. Cordero C. 2001 A different look at the false head of butterflies. Ecological Entomology 26, 106–108. (doi:10.1046/j.1365-2311.2001.00287.x)

