## Supplementary: for "Correlated evolution of multiple traits gives butterflies a false head"

**Supplementary files**

Table S1. List of databases and sources from where we sourced butterfly images to build the false head data

| **Sl. no.** | **Database/ source** | **Website** |
| --- | --- | --- |
| 01 | Butterflies of America | https://www.butterfliesofamerica.com |
| 02 | iNaturalist | https://www.inaturalist.org (limited only to images categorised as ‘Research Grade’) |
| 03 | Global Biodiversity Information Facility | https://www.gbif.org |
| 04 | Barcode of Life Data System | https://www.boldsystems.org |
| 05 | Australian Butterflies | https://www.purvision.com |
| 06 | Bali Wildlife | https://baliwildlife.com |
| 07 | Insecta Pro | https://insecta.pro |
| 08 | Butterflies of India | https://www.ifoundbutterflies.org |
| 09 | eBiodiversity Estonia | https://elurikkus.ee |
| 10 | Butterflies of Bulgaria | https://www.bulgarialeps.com |
| 11 | Lepiforum | https://lepiforum.org |
| 12 | Biodiversity Heritage Library | https://www.biodiversitylibrary.org |
| 13 | Flickr | https://www.flickr.com |
| 14 | Coffs Harbour Butterfly House | https://lepidoptera.butterflyhouse.com.au |
| 15 | Butterflies and Moths of North America | https://www.butterfliesandmoths.org |
| 16 | Butterflies of France | https://www.butterfliesoffrance.com |

Table S2. List of all Markov (Mk) models (ER = equal rates, ARD = all rates different) for the five false head traits fitted with *fitMk* using *phytools* with two different root priors. Rows representing the best-fitting models based on the Akaike Information Criteria (AIC) score are highlighted in bold.

| **Trait** | **Mk model** | **Root prior** | **log-likelihood** | **No. of parameter(s)** | **AIC** |
| --- | --- | --- | --- | --- | --- |
| false antennae | fitzjohn | -313.375598 | 1 | 628.751197 | fitzjohn |
| **false antennae** | fitzjohn | -310.892467 | 2 | 625.784933 | fitzjohn |
| spot | fitzjohn | -339.75873 | 1 | 681.51746 | fitzjohn |
| **spot** | fitzjohn | -333.361841 | 2 | 670.723682 | fitzjohn |
| conspicuous colouration | fitzjohn | -365.870697 | 1 | 733.741393 | fitzjohn |
| **conspicuous colouration** | fitzjohn | -335.972553 | 2 | 675.945105 | fitzjohn |
| false head contour | fitzjohn | -200.97282 | 1 | 403.94564 | fitzjohn |
| **false head contour** | fitzjohn | -193.906387 | 2 | 391.812773 | fitzjohn |
| convergent lines | fitzjohn | -80.1315154 | 1 | 162.263031 | fitzjohn |
| **convergent lines** | **fitzjohn** | **-76.2881433** | **2** | **156.576287** | **fitzjohn** |
| false antennae | flat | -313.575687 | 1 | 629.151373 | flat |
| **false antennae** | flat | -311.381797 | 2 | 626.763593 | flat |
| spot | flat | -340.257908 | 1 | 682.515816 | flat |
| **spot** | flat | -333.632243 | 2 | 671.264487 | flat |
| conspicuous colouration | flat | -366.395199 | 1 | 734.790398 | flat |
| **conspicuous colouration** | flat | -336.025141 | 2 | 676.050283 | flat |
| false head contour | flat | -201.174807 | 1 | 404.349613 | flat |
| **false head contour** | flat | -194.523591 | 2 | 393.047182 | flat |
| convergent lines | flat | -80.8208671 | 1 | 163.641734 | flat |
| **convergent lines** | **flat** | **-76.359184** | **2** | **156.718368** | **flat** |

Table S3. List of best-fitting ARD models for both root priors for the false head traits with their probability of the absence (0) and presence (1) at the root (equal for models with flat root prior) and the total number of character changes averaged for 1000 simulations.

| **Trait** | **Mk models** | | **Root priors** | **State probability at the root node** | | **Total no. of character changes** | | |
| --- | --- | --- | --- | --- | --- | --- | --- | --- |
|  |  |  |  | **0** | **1** | **0 to 1** | | **1 to 0** |
| false antennae | | ARD | fitzjohn | 0.10 | 0.90 | 19.19 | 83.64 | |
| false antennae | | ARD | flat | 0.50 | 0.50 | 19.44 | 82.98 | |
| spot | | ARD | fitzjohn | 0.78 | 0.22 | 43.62 | 74.20 | |
| spot | | ARD | flat | 0.50 | 0.50 | 43.22 | 75.82 | |
| conspicuous colouration | | ARD | fitzjohn | 0.62 | 0.38 | 55.35 | 88.34 | |
| conspicuous colouration | | ARD | flat | 0.50 | 0.50 | 55.23 | 89.83 | |
| false head contour | | ARD | fitzjohn | 0.04 | 0.96 | 10.12 | 47.03 | |
| false head contour | | ARD | flat | 0.50 | 0.50 | 10.26 | 46.60 | |
| convergent lines | | ARD | fitzjohn | 0.65 | 0.35 | 15.40 | 12.11 | |
| convergent lines | | ARD | flat | 0.50 | 0.50 | 15.67 | 13.33 | |

Table S4. Evolutionary correlates of pairwise discrete false head traits resulted from corHMM analyses using the phylogenetic framework of Espeland et al. (2018). Log-likelihood (lnL), AIC and corrected AIC or AICc of each evolutionary model are listed in the columns with the respective model’s name as the title. Model fit was assessed using the AICc. Pairwise traits with correlated evolution models are highlighted in bold.

| **Trait** | **Independent** | | | **Hidden Markov Independent** | | | **Correlated** | | | **Hidden Markov Correlated** | | |
| --- | --- | --- | --- | --- | --- | --- | --- | --- | --- | --- | --- | --- |
|  | lnL | AIC | AICc | lnL | AIC | AICc | lnL | AIC | AICc | lnL | AIC | AICc |
| **False antenna** |  |  |  |  |  |  |  |  |  |  |  |  |
| **Spot** | -176.86 | 361.72 | 361.93 | -152.37 | 324.73 | 325.95 | **-151.87** | **319.74** | **320.53** | -150.78 | 337.55 | 341.5 |
| **Conspicuous colouration** | -158.64 | 325.27 | 325.49 | -140.05 | 300.09 | 301.31 | **-139.58** | **295.15** | **295.94** | -138.96 | 313.92 | 317.88 |
| **False head contour** | -187.86 | 383.71 | 383.92 | -155.28 | 330.56 | 331.78 | **-156.89** | **329.77** | **330.55** | -154.26 | 344.52 | 348.48 |
| Convergent lines | -113.48 | 234.96 | 235.18 | **-111.73** | **243.45** | **244.66** | -112.76 | 241.52 | 242.3 | -111.53 | 259.05 | 263 |
| **Spot** |  |  |  |  |  |  |  |  |  |  |  |  |
| **Conspicuous colouration** | -126.68 | 261.35 | 261.56 | -109.3 | 238.59 | 239.81 | **-108.49** | **232.98** | **233.77** | -106.57 | 249.14 | 253.09 |
| **False head contour** | -155.9 | 319.79 | 320 | -134.25 | 288.49 | 289.7 | **-134.06** | **284.12** | **284.9** | -130.42 | 296.83 | 300.78 |
| Convergent lines | **-80.47** | **168.93** | **169.14** | -79.41 | 178.81 | 180.03 | -79.7 | 175.4 | 176.19 | -78.58 | 193.15 | 197.11 |
| **Conspicuous colouration** |  |  |  |  |  |  |  |  |  |  |  |  |
| **False head contour** | -137.67 | 283.34 | 283.56 | -125.06 | 270.12 | 271.34 | **-124.33** | **264.66** | **265.45** | -122.74 | 281.47 | 285.42 |
| Convergent lines | **-65.34** | **138.68** | **138.9** | -61.43 | 142.85 | 144.07 | -62.75 | 141.49 | 142.27 | -60.29 | 156.57 | 160.52 |
| **False head contour** |  |  |  |  |  |  |  |  |  |  |  |  |
| Convergent lines | **-91.8** | **191.59** | **191.8** | -91.12 | 202.23 | 203.44 | -91.08 | 198.15 | 198.94 | -90.33 | 216.65 | 220.61 |

Figure S1. Evolutionary transition rates between character states of all false head traits derived from the best-fitting ARD Mk models for flat root priors. 0 represents the absence of the trait, whereas 1 represents the presence. For each trait, the width of the arrows corresponds to the associated transition rate.

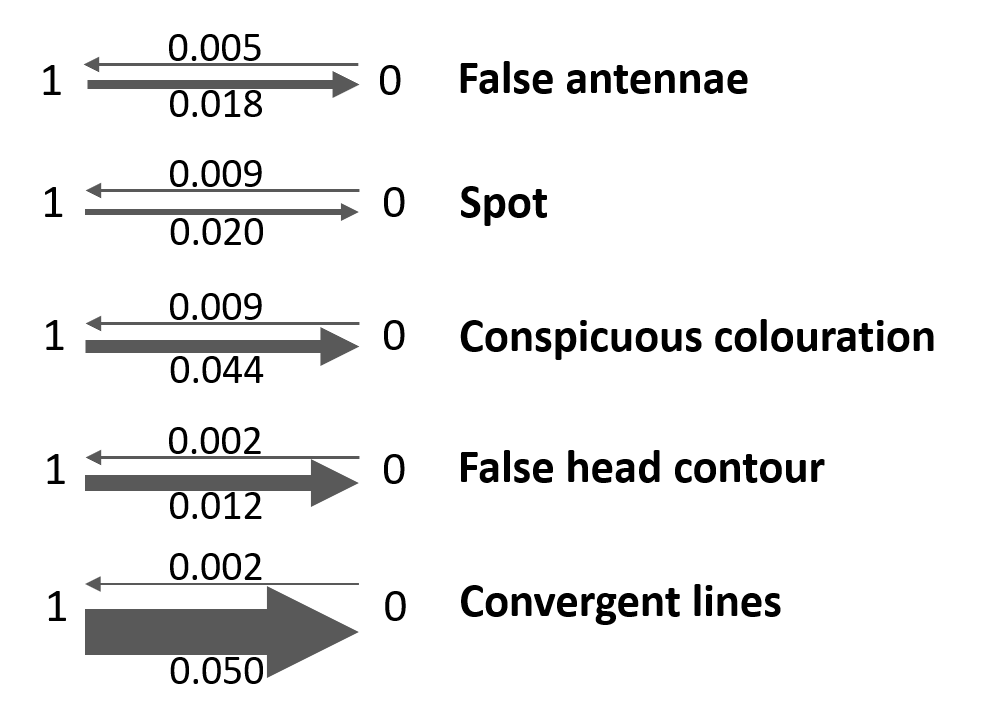

Figure S2. Replicated phylogenetic trees based on Espeland et al. (2018) with mapped false head traits at the side of the tip. The grey on the tip represents that the tip does not have the false head trait, while the presence of the trait is represented with coloured tips: blue (false antennae), dark green (spot), light blue (conspicuous colouration), yellow (false head contour) and convergent lines (red). All the pairwise combinations of the false head traits, with the exception of convergent lines, showed correlated evolution (Table S4).

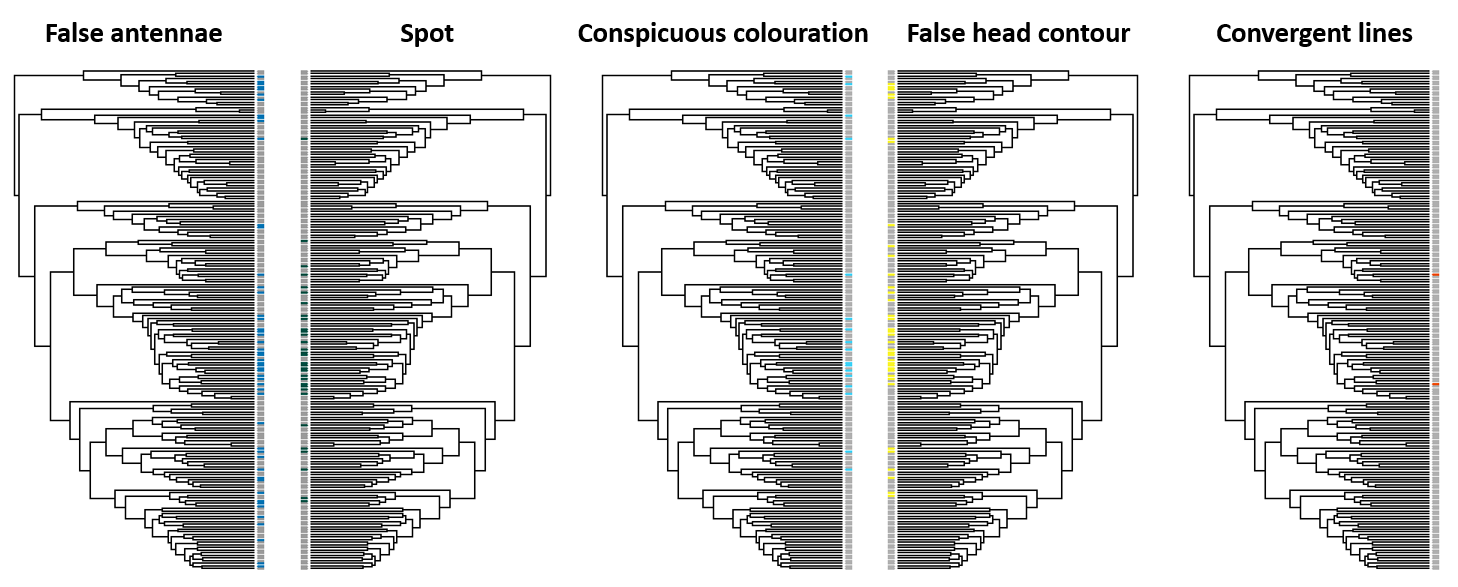

**Reference**

Espeland M *et al.* 2018 A Comprehensive and Dated Phylogenomic Analysis of Butterflies. *Current Biology* **28**, 770-778.e5. (doi:10.1016/j.cub.2018.01.061)
